# Evolution of sensory systems underlies the emergence of predatory feeding behaviours in nematodes

**DOI:** 10.1101/2025.03.24.644997

**Authors:** Marianne Roca, Güniz Göze Eren, Leonard Böger, Olena Didenko, Wen-sui Lo, Monika Scholz, James W. Lightfoot

## Abstract

Understanding how animal behaviour evolves remains a major challenge, with few studies linking genetic changes to differences in neural function and behaviour across species. Here, we identify specific sensory adaptations associated with the emergence of predatory feeding behaviours in the nematode *Pristionchus pacificus*. While *Caenorhabditis elegans* uses contact-dependent sensing primarily to avoid threats, *Pristionchus pacificus* has co-opted this modality to support both avoidance and prey detection, enabling context-dependent predatory behaviour. To uncover a potential mechanism underlying the evolution of *P. pacificus* prey perception, we mutated 27 canonical mechanosensory genes and assessed their function using behavioural assays, automated behavioural tracking, and a machine learning analysis of behavioural states. While several mutants showed mechanosensory defects, *Ppa-mec-6* mutants specifically also impaired prey detection, indicating the emergence of a novel mechanosensory module linked to predatory behaviour. Furthermore, disrupting both mechanosensation alongside chemosensation revealed a synergistic influence for these modalities. Crucially, both mechanosensation and chemosensation pathways converge in the same environmentally exposed IL2 neurons, and silencing these cells induced severe predation defects validating their importance for prey sensing. Thus, predation evolved through the co-option of mechanosensory and chemosensory systems that act together to shape the evolution of complex behavioural traits.

## Introduction

Sensory systems represent the primary mechanism through which an organism acquires environmental information. These signals can be used for finding food, avoiding predators, detecting mates, and navigating through complex surroundings. Detecting these environmental cues is dependent on specialised sensory systems which are often finely tuned to specific stimuli. Crucially, these systems are not static and their modification can profoundly affect the evolutionary trajectory of a species (1). Mechanisms of sensory system evolution include modifications to receptors, such as changes to receptor selectivity or sensitivity, as well as associated gene losses or gains (1). Examples of such events have been observed in several invertebrate species with changes in receptor function influencing feeding ecology in fruit flies (2), mosquitoes (3), and cockroaches (4). Similarly, in vertebrates, changes to sweet receptors have modified the feeding behaviours of songbirds and bats (5, 6) while in deep sea fish, a greatly expanded number of opsins has evolved to maximise their visual sensitivity in minimal light environments (7). Thus, sensory system evolution shapes the behavioural repertoire of an organism and facilitates its adaptation to its ecological niche.

In nematodes, a remarkable range of behaviours has evolved in accordance with the diverse ecologies found across the phyla. In particular, with their exceptional array of genetic and molecular tools, (8–10) alongside their distinct feeding ecologies, *Caenorhabditis elegans* and *Pristionchus pacificus* represent a powerful interspecies system for understanding the evolutionary adaptations influencing sensory perception. More specifically, while *C. elegans* is a microbial feeder, *P. pacificus* is an omnivorous species and not only feeds on bacteria but is also a predator of other nematodes (11). This expanded feeding behaviour is linked to a mouth-form plasticity with *P. pacificus* nematodes capable of developing one of two morphs. These are either the stenotomatous (st) or eurystomatous (eu) form. The st mouth is narrow with a single tooth and only permits bacterial feeding or scavenging, whereas the eu mouth is wider and possesses an additional tooth which facilitates both bacterial feeding and predation (11). Furthermore, predatory behaviour is directed towards not only other nematode species but also con-specifics. However, closely related strains are spared due to the presence of a robust kin-recognition system (12, 13). Crucially, how sensory systems evolved to accommodate the predatory behaviours is an open question.

In *C. elegans*, bacterial food is detected through well described foraging behaviour using their chemosensory and texture sensing systems (14, 15). In *P. pacificus*, little is known of its food and prey sensing mechanisms, however, they show specific sensory adaptations relevant to their ecology. In particular, *P. pacificus* are attracted to many bacterial species found concurrently in their environment and are also attracted to specific insect pheromones as *Pristionchus* are frequently found associated with scarab beetles (16). During food sensing and foraging, *P. pacificus* transitions between the docile bacterial feeding and aggressive predatory feeding states through the internal balance of octopamine and tyramine. These neuromodulators act on a group of head sensory neurons expressing the corresponding receptors (17). Contact between the nose of the predator and the surface of the prey during aggressive predatory bouts can initiate the activity of their movable tooth to puncture through the cuticle of the prey (11, 18, 19). However, the precise stimuli responsible for detecting prey contacts are currently unknown.

Here, to understand how the *P. pacificus* sensory systems have contributed to the emergence of predatory feeding we have investigated the role of mechanosensory adaptations for these behaviours. Using a candidate-based approach, we identified a crucial role for mechanosensation in predation. We find *Ppa-mec-6* is necessary for this process and may contribute to these behaviours through a previously undescribed predatory-associated mechanosensory channel. Furthermore, we demonstrate that mechanosensation is required alongside chemosensory inputs and that these acts synergistically to execute efficient predation behaviours. We also reveal that chemosensory and mechanosensory components are expressed in the same externally projecting and environmentally exposed neurons, indicating functional convergence into the same sensory circuits. Finally, we show that the functional inhibition of these neurons indeed disrupts predation.

## Results

### Mechanosensation is involved in prey detection

*P. pacificus* is a voracious predator of other nematode larvae including *C. elegans* and this feeding behaviour requires contact between predator and prey (Fig. 1A). Accordingly, we investigated the importance of *P. pacificus* mechanosensation for detecting prey and a potential role in the evolution of its predation behaviours. *Cel-mec-3* is critical transcription factor involved in the development and function of mechanosensory neurons in *C. elegans*, and animals with mutations in this gene fail to respond to tactile stimuli (20). In *P. pacificus*, using a *Ppa-mec-3* transcriptional reporter, we observed expression in several neurons with similar cell body placements to the *C. elegans* mechanosensory neurons FLP, AVM, ALM, PVM, PVD, and PLM (Fig. 1B), (20–22). To assess its role in mechanosensation in *P. pacificus*, we generated a mutation in the *Ppa-mec-3* orthologue using CRISPR/Cas9. Similar to *C. elegans*, we found *Ppa-mec-3* mutants fail to react to several classic mechanosensory assays including harsh touch, gentle touch, and nose touch assays (Fig. 1C-E). Taken together, this indicates a conserved role for *Ppa-mec-3* between species. Next, to assess a potential role for *Ppa-mec-3* in predatory feeding, we performed previously established corpse assays (11). In corpse assays, starved *P. pacificus* adults are placed on a plate saturated with *C. elegans* larval prey and predation success is determined by the number of corpses after a designated time interval (supplementary Fig. 1A). Here, *Ppa-mec-3* predators show a significant reduction in killing ability, with fewer larval corpses generated by predators than observed in control assays with wildtype predators (Fig 1F). Thus, mechanosensory systems have acquired additional functions in *P. pacificus* and are necessary for efficient prey detection.

**Figure 1.**
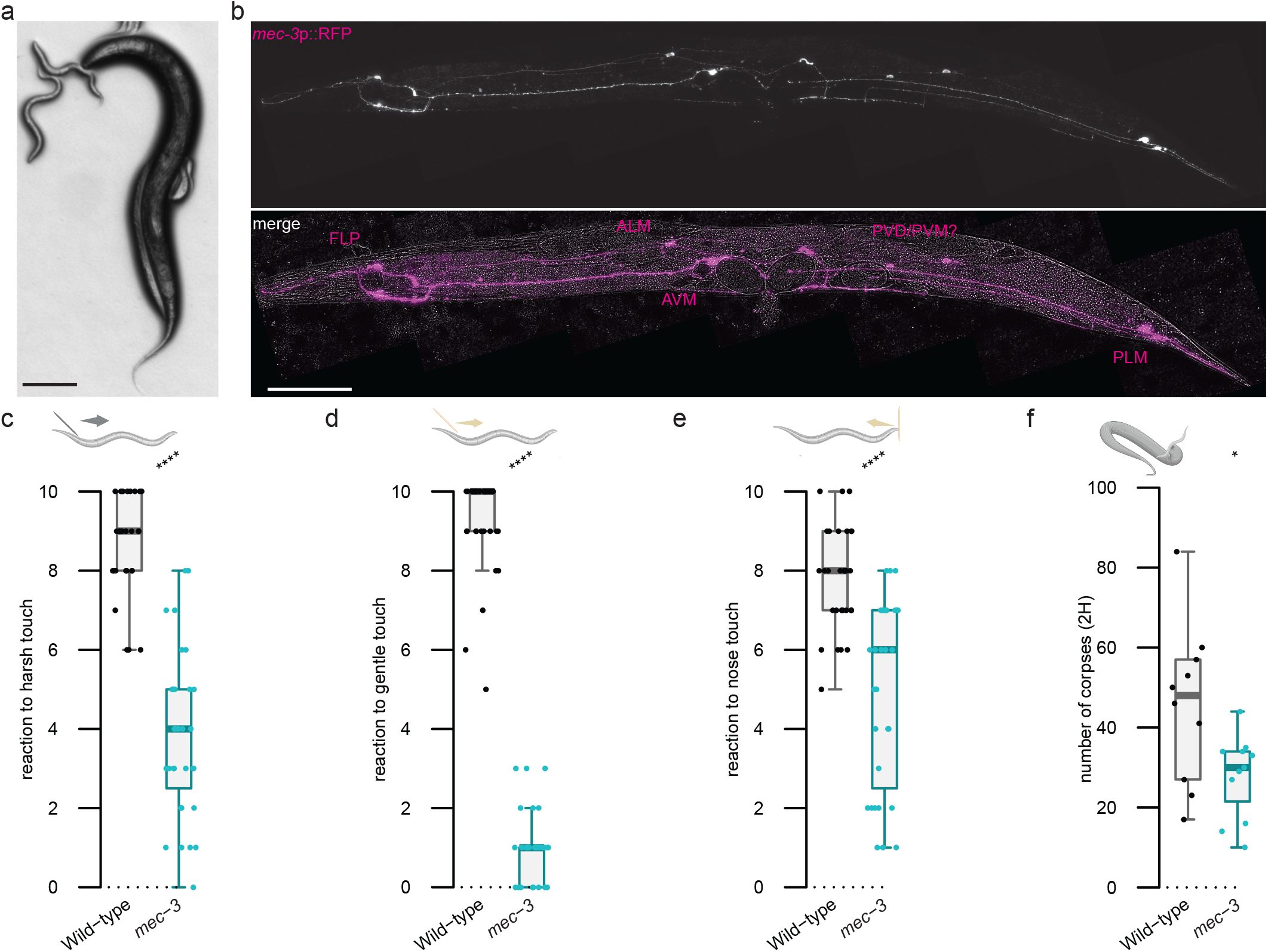
*Ppa-mec-3* regulates touch avoidance and prey detection. (A) A *P. pacificus* adult predator (right) attacks a *C. elegans* larvae (left). Scale bar is 100 μm. (B) Representative image of a *P. pacificus* adult expressing *Ppa-mec-3p*::RFP (top, magenta). Putative *P. pacificus* neuronal identity is based on *C. elegans* soma placement and known mechanosensory function. Scale bar is 100μm. (C) Mechanosensory assays to harsh touch, (D) gentle touch, and (E) nose touch. Each assessment is the result of ten consecutive trials of each worm. At least 30 worms were tested per strain. (F) Number of *C. elegans* corpses counted after two hours of contact with the indicated *P. pacificus* strains as predator. At least 10 assays were performed. Statistical tests: one-direction Wilcoxon Mann Whitney with Benjamini-Hochberg correction, non-significant (ns), p-value ≤ 0.05 (*), ≤ 0.01 (**), ≤ 0.001 (***), ≤ 0.0001 (****). Schematics were made with biorender.

### Mechanosensory genes have diversified across evolution

Having established the importance of mechanosensation for predation, we next explored the conservation of mechanosensory gene networks across nematode evolution. Frequent gene losses and gains are observed between nematode species and this can be associated with novel evolutionary processes (23). Therefore, we initially conducted a bioinformatic study of the mechanosensory systems across 8 nematode species to investigate the evolution of this sensory modality across the nematode phyla (Fig. 2A). Our analysis consisted of the two free-living species *C. elegans* and *P. pacificus* belonging to clade 5 as well as several obligate parasite species of both plants and animals (23). We included orthologues of the *C. elegans* predicted mechanostransduction channels belonging to the degenerin–epithelial Na+ channels (DEG/ENaC) and transient receptor potential (TRP) gene families (22). We also included other known mechanosensory genes such as *mec-6*, a paraoxonase-like protein which acts as a chaperone to ensure channel function (24), and *pezo-1*, due to its reported role in food sensation (25). Compared to free-living nematodes, parasitic nematodes frequently had a lower number of genes encoding mechanotransduction channels, perhaps indicative of adaptations to their specific lifestyle. In contrast, comparisons between the free-living nematodes *C. elegans* and *P. pacificus* generally showed a similar representation of gene families with a few exceptions (Fig. 2A). These include the four *trpl* genes which are specific to *C. elegans* but absent in other species including *P. pacificus* (22). Additionally, there are two large gene expansions in *P. pacificus* among the DEG/ENaC gene family with five copies of *deg-1* in *P. pacificus* compared to one in *C. elegans* and 11 *egas* genes in *P. pacificus* compared to four in *C. elegans*. Little is known regarding the function of *egas* in *C. elegans* and many of the *P. pacificus egas* paralogues lack RNA-seq data to validate their expression (10). However, *deg-1* has been shown in *C. elegans* to be required for the induction of mechanoreceptor current in the sensory ASH neuron (26). Therefore, mechanosensory gene networks appear relatively stable between *C. elegans* and *P. pacificus* but are more stochastic in parasitic nematodes with their more complex life-styles and host interactions.

**Figure 2.**
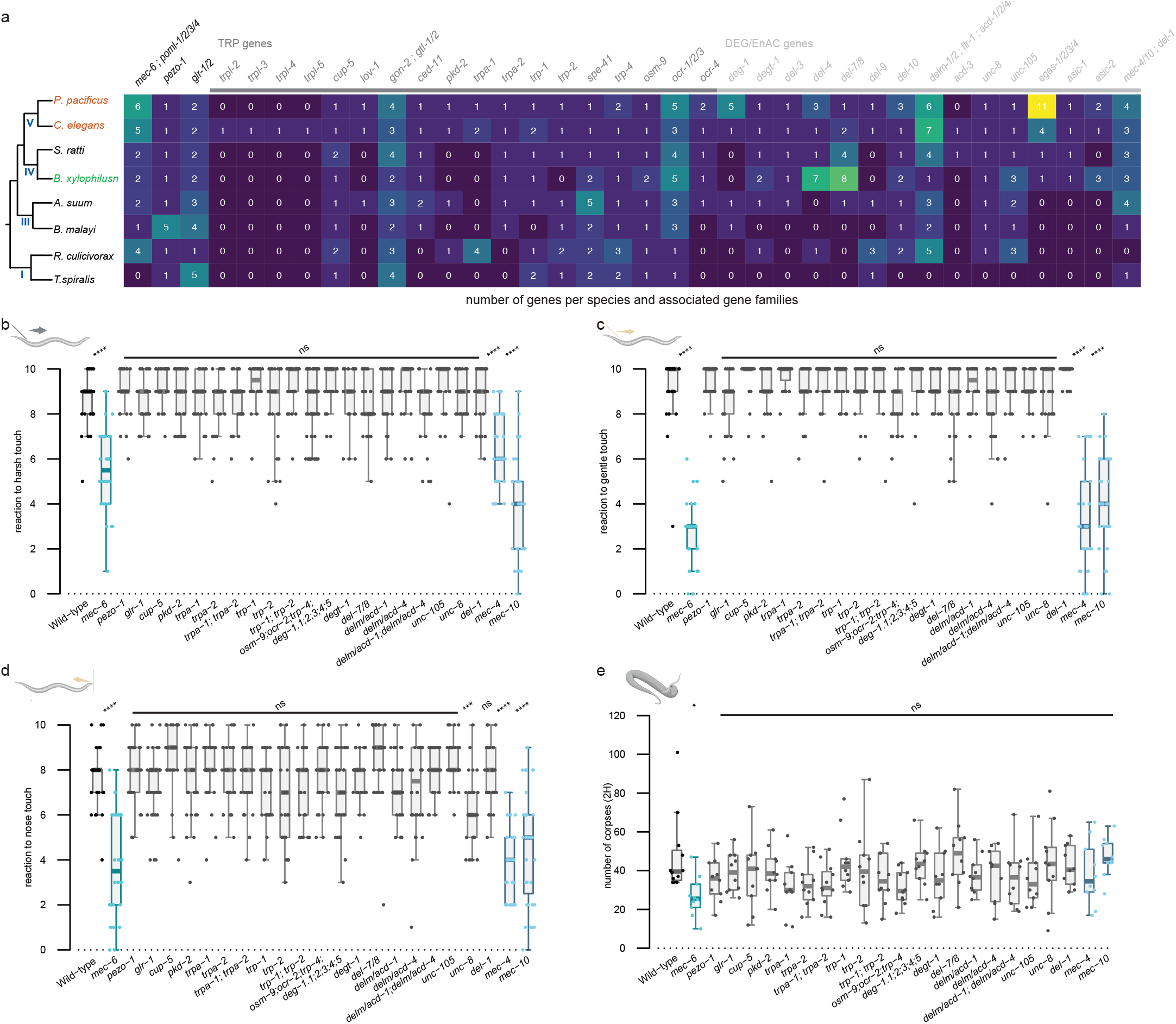
Mechanosensation is required for efficient predation. (A) Table analysing the number of mechanosensory gene paralogues compared to *C. elegans*. Species of free-living nematodes (brown), plant parasite nematodes (green) and animal parasite nematodes (black) are compared. Their evolutionary tree is presented on the left with the nematode clade marked. (B) Mechanosensory assays to harsh touch, (C) gentle touch, and (D) nose touch for *P. pacificus* mutants in putative mechanosensory genes. Each assessment is the result of ten consecutive trials of each worm. At least 30 worms were tested per strain. (E) Number of *C. elegans* corpses counted after two hours of contact with the indicated *P. pacificus* putative mechanosensory mutants as predators. At least 10 assays were performed per mutant. Statistical tests: one direction Wilcoxon Mann Whitney with Benjamini-Hochberg correction, non-significant (ns), p-value ≤ 0.05 (*), ≤ 0.01 (**), ≤ 0.001 (***), ≤ 0.0001 (****). Schematics were made with biorender.

With the identification of genes potentially acting as mechanosensory channels in *P. pacificus*, we conducted a candidate-based approach to identify components necessary for efficient prey detection through *P. pacificus* nose contact. Candidates were selected based on known mechanosensory phenotypes in *C. elegans* together with further validation through available *P. pacificus* expression data (10, 27). Accordingly, we selected genes to target by CRISPR/Cas9 which included both TRP and DEG/ENaC families as well as other associated mechanosensory genes. We selected the DEG/ENaC channels *Ppa-mec-4* and *Ppa-mec-10* as well as *Ppa-mec-6*. In *C. elegans, Cel-mec-4* and *Cel-mec-10* are both involved in the gentle touch response (22, 28) and *Cel-mec-6* has been shown to directly interact with *Cel-mec-4* and *Cel-mec-10* to aid the function and stability of this mechanosensitive channel (24, 29). We targeted *Ppa-osm-9, Ppa-ocr-2* and *Ppa-trp-4* together as in *C. elegans* these genes have an additive effect reducing its nose touch responsiveness (30). Only *P. pacificus* triple mutants for these genes were analysed. *C. elegans* nose touch response also requires *Cel-delm-1* and *Cel-delm-2* which are expressed in glial cells (31). However, we could not identify one-to-one orthologues for these two genes in *P. pacificus* as both paralogous sequences are closer to each other than to either *Cel-delm* or *Cel-acd* genes. Therefore, these were named *delm/acd* and we mutated both candidates in *P. pacificus*. In *C. elegans, Cel-del-7* and *Cel-del-8* are DEG/ENaC ion channel family members, with the former involved in mechanotransduction (32). However, there is only one paralogue in *P. pacificus*, which we mutated and named *Ppa-del-7/8*. We also generated a quintuple mutant of all paralogues of DEG/ENaC *Ppa-deg-1*, and targeted the DEG/ENaC potential proprioception genes *Ppa-del-1, Ppa-unc-8* and *Ppa-unc-105* (22, 33). To further represent the TRP gene family, we included *Ppa-trp-1, Ppa-trp-2, Ppa-pkd-2, Ppa-cup-5* and *Ppa-trpa-1, which*, in *C. elegans*, are expressed in many neurons and most have roles in mechanotransduction (30, 31, 34, 35). Finally, we targeted the ionotropic glutamate receptor *Ppa-glr-1* due to its role in mediating excitatory neurotransmission in touch receptor neurons (36), and *Ppa-pezo-1* due to potential roles in food sensing (25). This resulted in 26 genes of interest, and we investigated two alleles for each of these genes, which were predicted to cause loss-of-function mutations (Supplementary Table 1).

### MEC-6 is required for efficient *P. pacificus* predation

Initially, we tested all of our mutants for mechanosensory defects using the three touch assays adapted from previous *C. elegans* studies. These assays measure the animal response to harsh touch, gentle touch, and nose touch (22). The majority of the *P. pacificus* mutants did not show mechanosensory deficient phenotypes. However, *Ppa-mec-4, Ppa-mec-10*, and *Ppa-mec-6* showed mechanosensory defects similar to those described in *C. elegans*, and also similar to those observed in *Ppa-mec-3*, resulting in a response to touch only half the time or less (Fig. 2B-D and supplementary Fig. 1B-D). Furthermore, all four mutants were also more lethargic than wildtype worms, phenocopying *C. elegans* mechanosensory mutants (28). Thus, as in *C. elegans, Ppa-mec-4, Ppa-mec-10, Ppa-mec-3*, and *Ppa-mec-6* are required for touch sensation in *P. pacificus*. While previous reports in *C. elegans* have not identified a role for harsh touch sensation in *Cel-mec-4* (30), this discrepancy between the two species may stem from subtle species-specific sensitivity differences. Next, we tested all of our mutants for defects in predation using the previously described corpse assays. While the majority of mutants displayed wildtype levels of predation, reduced killing was observed in *Ppa-mec-6* (Fig 2E and supplementary Fig. 1E). This is similar to the defect observed in the *Ppa-mec-3* mutants (Fig. 1F). Importantly, no predatory defect was observed in *Ppa-mec-4* or *Ppa-mec-10*, indicating that the predatory abnormality is not the consequence of a general mechanosensory defect, but rather is specific to the function of *Ppa-mec-3* and *Ppa-mec-6*. In *C. elegans, Cel-mec-6* is involved in the assembly and function of mechanosensory ion channels and interacts with *Cel-mec-4* and *Cel-mec-10* to form ion channel complexes that are essential for mechanotransduction (24, 29). Our data indicates that while the *Ppa-mec-4* and *Ppa-mec-10* channels are not required for prey detection in *P. pacificus, Ppa-mec-6* may function as part of an additional, as yet unknown, mechanosensory ion channel involved in detecting prey contact. Thus, specific mechanosensory inputs are necessary for prey detection in *P. pacificus*. However, as predation is not fully abolished, additional sensory inputs also contribute to prey detection ability.

### Mechanosensation and other predatory-associated traits

Predation in *P. pacificus* is dependent on the formation of the eu mouth morph which is determined by a multitude of environmental and genetic factors (37, 38). To assess if mechanosensation also influences the mouth morph fate, we screened our mutant library for any effect on mouth form ratio. All mutants showed wildtype levels of eu to st morphs (supplementary Fig. 2A and B). In addition, *P. pacificus* has a robust kin-recognition mechanism which prevents attacks on its own progeny and close relatives, while facilitating the cannibalism of other con-specific competitors (12, 13). Due to the prey detection defect observed in *Ppa-mec-3* and *Ppa-mec-6* we also tested these mutants for kin-recognition defects. For both *Ppa-mec-3* and *Ppa-mec-6* mutants, kin-recognition was robustly maintained indicating no deficiency in this process (supplementary Fig. 2C). Therefore, defects in mechanosensation are specific to prey detection and do not influence other predatory traits.

### Chemosensation and mechanosensation synergistically influence prey detection

The partial defect in prey detection observed in the *Ppa-mec-3* and *Ppa-mec-6* mutants is similar to that previously describe in cilia deficient mutants (18). Cilia are complex organelles, and in nematodes they are essential for the detection of many environmental cues. In both *C. elegans* and *P. pacificus*, some of the strongest cilia deficiencies are observed in mutants defective for the transcription factor, DAF-19, which acts as the master regulator of ciliogenesis (18, 39, 40). Curiously, while mutations in *Cel-daf-19* cause aberrant dauer entry, no abnormal dauer phenotypes result from *Ppa-daf-19* mutations in *P. pacificus*, reinforcing the distinct evolutionary trajectories regulating this developmental process between these species (41). Furthermore, accompanying the numerous chemosensory defects in *Ppa-daf-19*, these mutants are also defective in prey detection (18). Therefore, to disentangle the roles of chemosensation and mechanosensation, we assessed the *Ppa-mec-3* and *Ppa-mec-6* mutants alongside *Ppa-daf-19*. We found that while *Ppa-daf-19* mutants showed chemosensory defects in their attraction to a bacterial food source and in their aversion to octanol, both *Ppa-mec-3* and *Ppa-mec-6* responded as wildtype animals to these cues (Fig. 3 A and B). This indicates both *Ppa-mec-3* and *Ppa-mec-6* are not defective for chemosensation. In contrast, while both *Ppa-mec-3* and *Ppa-mec-6* are mechanosensory defective, the touch response of *Ppa-daf-19* mutants was similar to wild type animals (supplementary Fig. 3 B-D). A similar observation has also been reported in *C. elegans*, where only mild mechanosensory defects are observed in *Cel-daf-19* mutants (42). Thus, mechanosensory defects appear specific to *Ppa-mec-3* and *Ppa-mec-6*, while *Ppa-daf-* 19 mutants are mostly associated with chemosensory defects. As both groups of mutants are defective for prey detection, but neither fully abrogates the *P. pacificus* predatory abilities, we next assessed if these sensory inputs act synergistically. We found that the predation defect was further exacerbated in double mutants of *Ppa-mec-6; Ppa-daf-19* as well as *Ppa-mec-3; Ppa-mec-6; Ppa-daf-19* triple mutants, but crucially, some killing was still maintained (Fig. 3C). Therefore, chemosensation and mechanosensation act together for prey detection. Remaining killing abilities may act through some residual function in these sensory pathways or through another as yet unidentified stimuli.

**Figure 3:**
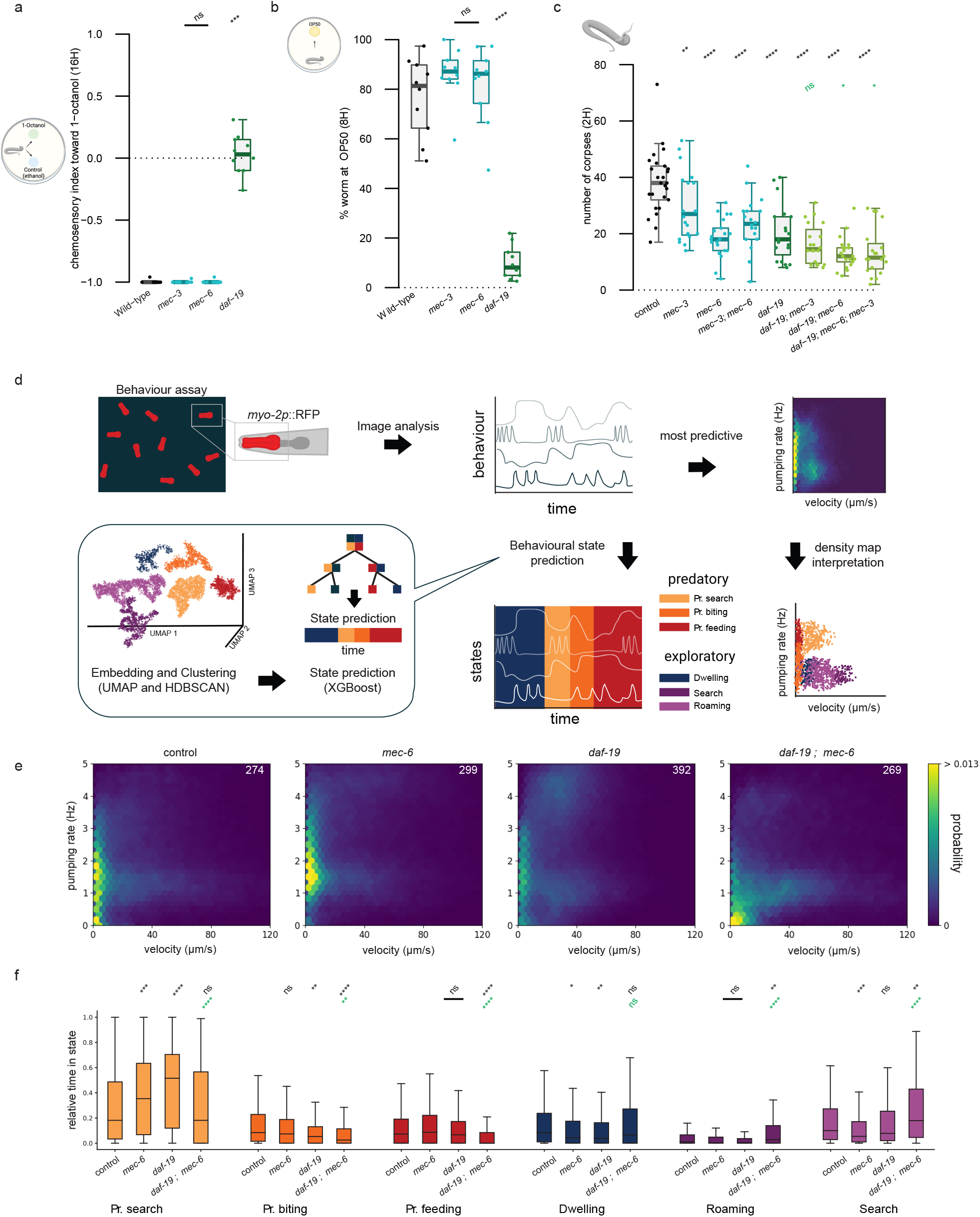
Chemosensation and mechanosensation synergically influence prey detection. (A) Chemotaxis index analysing the aversive response of *Ppa-mec-3, Ppa-mec-6* and *Ppa-daf-19* to 1-octanol. Each genotype was assessed 10 time. (B) Percentage of *Ppa-mec-3, Ppa-mec-6* and *Ppa-daf-19* mutants finding a bacterial OP50 food source after 8 hours. Each genotype was assessed 10 times. (C) Number of *C. elegans* corpses generated after two hours of exposure to the indicated *P. pacificus* mutants as predator. At least 20 assays were performed. (D) Schematic of behavioural tracking and machine learning workflow to track feeding behaviour and determine behavioural states based on a *Ppa-myo-2*::RFP pharyngeal fluorescence marker (established in Eren et al 2024). (E) Joint probability density map of velocity (μm/s) and pumping rate (Hz) for animals corresponding to the genotypes *Ppa-mec-6, Ppa-daf-19*, and *Ppa-mec-6; Ppa-daf-19*. The number of worms is indicated in the top right corner. (F) Time spent in each behavioural state normalised to the total track duration. Statistical tests: Significance from comparison to wild-type (black) and to *daf-19* single mutant (green) was assessed using one direction Wilcoxon Mann Whitney with Benjamini-Hochberg correction (A-C), Mann-Whitney U-test with a Bonferroni correction (F). non-significant (ns), p-value ≤ 0.05 (*), ≤ 0.01 (**), ≤ 0.001 (***), ≤ 0.0001 (****). Schematics were made with biorender. Predatory (Pr.)

### Behavioural tracking and state predictions

Quantifying the number of *C. elegans* corpses generated by *P. pacificus* predators provides a robust assessment of overall killing ability (11). However, understanding the sensory-motor transformations underpinning prey contact requires a more detailed behavioural analysis. We recently developed an automated behavioural tracking and machine learning model to identify and quantify aggressive behavioural states associated with predation and further analyse these complex behaviours (17, 43). Using this method, the *P. pacificus* behavioural repertoire can be divided into six states. These consist of three predatory associated states including ‘predatory search’, ‘predatory biting’ and ‘predatory feeding’ (Fig. 3D). Additionally, three non-predatory behaviours including a ‘dwelling’ state and two ‘roaming’ states exist which are similar to behavioural states observed in *C. elegans* (15, 17). The two most predictive features of predation states are velocity and pumping rate. By observing the joint distribution of these two features, it is possible to visualize the prevalence of predatory and non-predatory states (Fig. 3D). We used a machine learning model that was previously trained on behavioural data using both unsupervised and supervised methods (17) to investigate behavioural state occupancy in our predation mutants compared to wildtype animals (Fig. 3D-F, and supplementary Fig. 3E-G). In the mechanosensory defective *Ppa-mec-6* and *Ppa-mec-3*, as well as the chemosensory defective *Ppa-daf-19* mutants, we found a significant increase in the duration and total time spent into the ‘predatory search’ state. This may be indicative of a compensatory mechanism whereby the loss of one sense enhances utilisation of the remaining senses. This is further supported by the absence of the increased ‘predatory search’ state in mutants lacking both sensory modalities (Fig. 3F). Instead, these *Ppa-mec-6; Ppa-daf-19* double mutants show a significant increase in the non-predatory ‘search’ and ‘roaming’ states, indicating a switch away from predatory states and into exploratory behaviours. This is similar to observations in *C. elegans* where the loss of one or multiple sensory modalities results in modifications to the performance of the remaining senses and induces behavioural adjustments (28). Furthermore, ‘predatory biting’ state occupancy was similar to wildtype animals in both *Ppa-mec-6* and *Ppa-mec-3* mutants while, in *Ppa-daf-19* this state was significantly reduced. Therefore, predatory behavioural states are modulated through signals received through the *P. pacificus* sensory systems with both chemosensory and mechanosensory modalities influencing distinct but overlapping aspects of prey detection.

### Prey detection systems converge on the IL2 sensory neurons

As distinct mechanosensory and chemosensory inputs contribute to the *P. pacificus* predatory abilities, we investigated if they act through discrete independent neurons, or if instead, they converge into the same sensory pathways. To resolve this, we generated a reporter strain expressing *Ppa-mec-6p*::Venus and *Ppa-daf-19p*::RFP and identified the specific prey detection neurons involved. *Ppa-mec-6p*::Venus and *Ppa-daf-19p*::RFP co-localised and were robustly expressed in the *P. pacificus* IL2 neurons (Fig. 4A and B). These were identified based on soma position and their unique morphological features (44). Importantly, the anterior processes from this group of six neurons project into the external environment through exposed sensory endings making them strong candidates for prey detection (Fig. 4C). Additionally, we observed that *Ppa-mec-6* is co-expressed with *Ppa-mec-3* in the putative *P. pacificus* ALM and PLM body neurons as well as the FLP head neurons (supplementary Fig. 4A and B). Alongside this, *Ppa-daf-19* shows low expression in few other head neurons (Fig. 4B). As in *C. elegans*, many sensory neurons in *P. pacificus* are ciliated and this is not restricted to the IL2 sensory neurons (40). Therefore, it is somewhat surprising that the most robust *Ppa-daf-19* expression is observed in the IL2 neuron cluster. However, as there are several large introns in the *Ppa-daf-19* gene structure which contain enhancers and other epigenetic signatures (45), we predict these may contribute additional regulatory elements for further expression in other ciliated neurons. These may currently be missing from our *Ppa-daf-19p*::RFP reporter. Thus, both mechanosensory and chemosensory prey detection systems converge on the same head neurons with sensory projections exposed to the external environment, likely acting as the first point of contact with potential prey.

**Figure 4:**
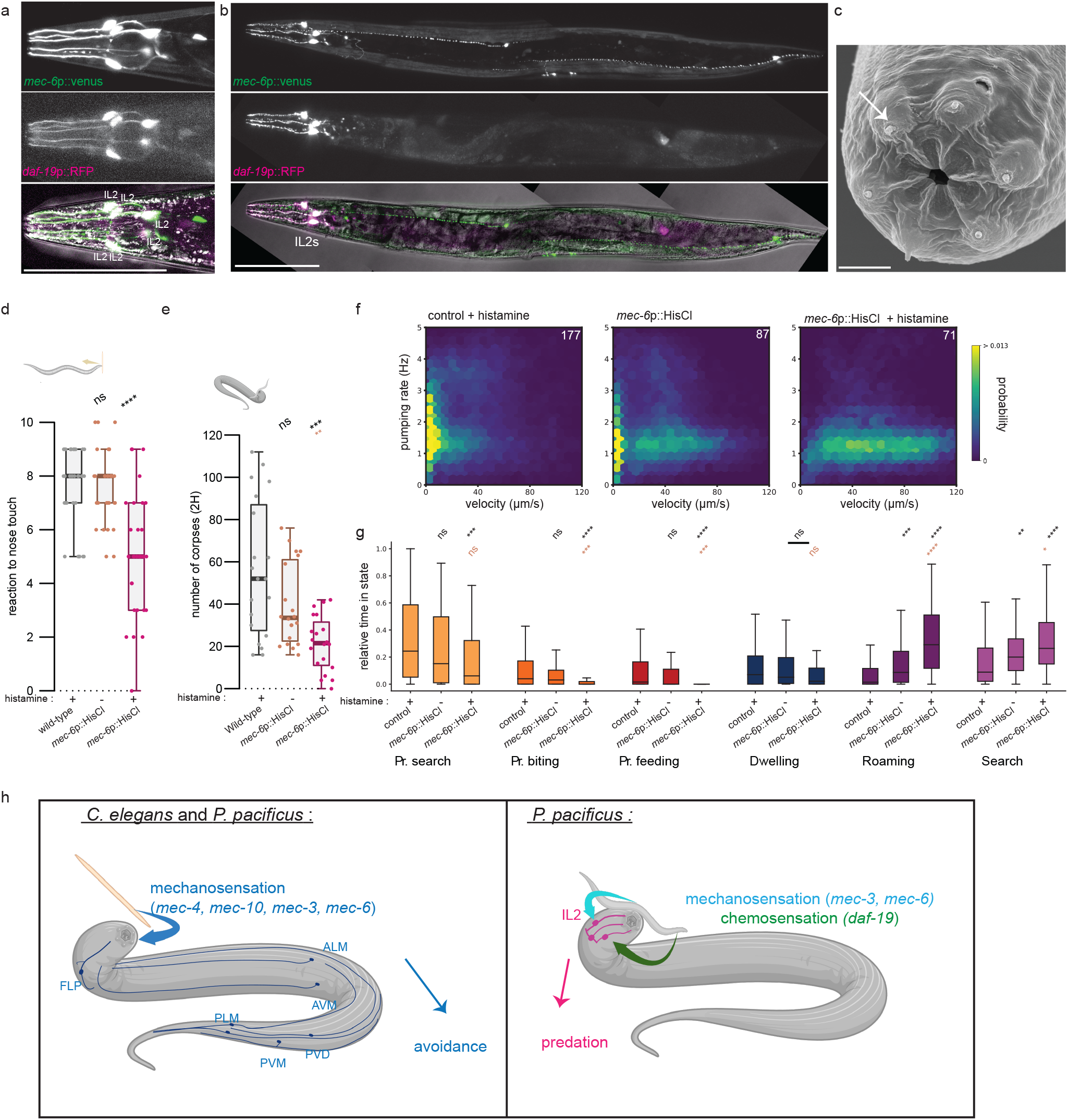
Predation is disrupted by inhibition of *Ppa-mec-6* expressing cells. (A) Head of a worm expressing *mec-6p*::Venus (top, green) and *daf-19p*::RFP (middle, magenta). Co-expression of both *Ppa-mec-6* and *Ppa-daf-19* visible in the merge image. IL2 neurons are indicated. Scale bar is 100 μm. (B) Representative image of a worms expressing *Ppa-mec-6p*::Venus (top, green) and *Ppa-daf-19p*::RFP (middle, magenta) and the merge (bottom). Scale bar is 100 μm. (C) Scanning electron microscopy image of the anterior of a *P. pacificus* adult. The *P. pacificus* mouth opening is surrounded by protrusions of the 6 inner labial IL2 sensilla neurons (arrow). Scale bar is 3 μm (D) Mechanosensory assays of nose touch for *P. pacificus* worms for each condition: wild-type worms treated with histamine and worms expressing HisCl channel under the control of *Ppa-mec-6* promotor in absence (-) or presence (+) of histamine. Each datapoint is the result of ten consecutive trials on an individual worm. At least 30 worms were tested per condition. (E) Number of *C. elegans* corpses generated after two hours of exposure to the indicated *P. pacificus* worms as predator in absence (-) or presence (+) of histamine. 20 assays were performed. (F) Joint probability density map of velocity (μm/s) and pumping rate (Hz) for animals corresponding to condition described in (D). The number of worms is indicated in the top right corner. (G) Time spent in each behavioural state normalised to the total track duration. (H) Putative model of the evolutionary diverse role for mechanosensation between *C. elegans* and *P. pacificus*. In both species, mechanosensation triggers an avoidance response which rely on *mec-3, mec-6, mec-4* and *mec-10*. In addition, in *P. pacificus*, contact can trigger predatory behaviour when the IL2 perceive mechanical or chemical stimuli. The perception of the predatory specific mechanical stimuli, acts through the mechanosensory pathway components *Ppa-mec-3* and *Ppa-mec-6*. Statistical tests: Significance from comparison to wild-type (black) and to *Ppa-mec-6p*::HisCl untreated worms (brown) was assessed using one direction Wilcoxon Mann Whitney with Benjamini-Hochberg correction(D-E), Mann-Whitney U-test with a Bonferroni correction (G). non-significant (ns), p-value ≤ 0.05 (*), ≤ 0.01 (**), ≤ 0.001 (***), ≤ 0.0001 (****). Predatory (Pr.) Schematic was made with biorender.

### Functional inhibition of *Ppa-mec-6* expressing neurons disrupts mechanosensation and predation

Finally, with the convergence of sensory stimuli in the IL2 neurons and the importance of *Ppa-mec-6* for prey detection, we investigated the requirements of *Ppa-mec-6* expressing cells for predation. We expressed a histamine-gated chloride channel (HisCl) under the control of the *Ppa-mec-6* promoter. This method, recently established in *P. pacificus*, enables inducible inhibition of target neurons upon the exogenous addition of histamine as neurons become hyperpolarised (46). We first verified that the construct and treatment did not affect worm health. Developmental time was the same across wildtype and transgenic *Ppa-mec-6p*::HisCl animals, both alone and in the presence of histamine, indicating no adverse effects (supplementary Fig. 4C). Furthermore, mouth form also remained at wildtype ratios indicating *Ppa-mec-6* expressing cells, including the IL2 sensory neurons, do not influence this trait (supplementary Fig. 4D). Next, we assessed the requirement of the *Ppa-mec-6* expressing neurons for mechanosensation and chemosensation. Both body touch and nose touch response were significantly reduced upon silencing of the *Ppa-mec-6* positive neurons although the effect on nose touch was much greater than the body touch response (Fig. 4D, supplementary Fig. 4E and F). We also observed no defect in the response to our chemosensory stimuli upon neuronal silencing (supplementary Fig. 4G and H). Finally, we tested the consequences of this inhibition on predation. Both the number of corpses and prevalence of predatory states were reduced when *Ppa-mec-6p*::HisCl expressing worms were treated with histamine (Fig. 4E-G and supplementary Fig. 4I). Automated identification of the different behavioural states indicated that time dedicated to all three predatory behaviours - search, biting and feeding were reduced. These reductions were accompanied by an increase in two of the exploratory behaviours - search and roaming. Thus, neurons expressing *Ppa-mec-6*, including the IL2 sensory neurons, are required for predation.

## Discussion

A central aspect associated with the evolution of predatory behaviours hinges on an organism’s ability to detect potential prey. Fundamental to this are adaptations to sensory abilities that enable an organism to perceive, process and respond to potential prey cues and direct predatory behaviours appropriately. In the nematode *P. pacificus*, we have found that the evolution of predation is associated with the diversification and specialisation of both mechanosensory and chemosensory modalities to also encompass prey detection (Fig. 4H). This is beyond their previously described function in the model species *C. elegans*. The co-option and neofunctionalisation of sensory systems are a frequently observed phenomenon. For example, the lateral line system, used to detect water movements and vibrations in many fish species, has evolved electroreception capabilities in sharks, rays and skates (47). Similarly, certain snake species including pit vipers, pythons and some boas have evolved infra-red detection through adaptations to the temperature-sensitive skin thermotransduction receptor TRPA1. This enables these species to hunt warm blooded prey in near total darkness, while in other snakes it is a mechanism to regulate body temperature (48). Examples of sensory co-option events are not restricted to animals and can also be readily found in plants including carnivorous Venus flytrap and sundew species, whereby ancestral mechanosensory channels are instead used for sensing and prey capture (49). Therefore, our findings support a well-established link between sensory system evolution and behavioural diversity across a wide range of taxa.

In *P. pacificus*, the co-option and integration of multiple sensory modalities, suggests that predation is not solely dependent on enhancements and adaptations to a single sensory pathway but rather on the evolution of multiple interacting systems which fine tune prey capture. In this nematode species, it is both chemosensation and mechanosensation that are required for efficient predation. Similar multimodal mechanisms of predation can be observed in other invertebrates, including other species which also lack visual inputs or those which hunt in low light environments. These include species of octopus, which utilise chemotactile receptors on their tentacles to identify potential prey through a mechanism of contact-dependent aquatic chemosensation (50). Additionally, cone snails are carnivorous molluscs that hunt fish, worms or other snails and utilize chemosensation to first detect the presence of chemicals secreted by their prey, and subsequently use vibration sensing and mechanoreception to determine prey position. They then shoot a venomous harpoon-like tooth to immediately immobilise and feed on their target (51). In *P. pacificus*, our data shows that both chemosensation and mechanosensation are important for predation, and defects in either system significantly reduce killing ability. Additionally, from our behavioural state studies of these mutants, we find defects in *Ppa-daf-19* chemosensory mutants are more severe and are associated with aberrations in the ‘predatory biting’ state, alongside increased ‘predatory search’ occupancy. A similar ‘predatory search’ occupancy increase is also detected in *Ppa-mec-3* and *Ppa-mec-6* mutants. We hypothesize that the increased ‘predatory search’ state occupancy in both sensory modalities is a compensatory mechanism whereby *P. pacificus* switches dependency to the unimpaired sense in an attempt to maintain its predatory behaviours. This is further validated by the loss of this increase in mutants lacking both the predatory-associated mechanosensory and chemosensory components. Indeed, sensory loss in *C. elegans* has also been shown to alter the performance of remaining sensory modalities resulting in distinct behaviours (28). Furthermore, while the mechanosensory *Ppa-mec-6* and *Ppa-mec-3* as well as chemosensory *Ppa-daf-19* mutants show reduced killing abilities, *Ppa-mec-6* and *Ppa-mec-3* mutants were found to occupy ‘predatory biting’ states similar to wildtype, while this state was reduced in *Ppa-daf-19*. Therefore, both sensory inputs likely have specific functions during predation, such that chemosensation is potentially required throughout these behaviours while mechanosensory systems may be associated with the initial steps of the predator-prey contact.

Both *daf-19* and *mec-*3 are transcription factors and influence genes associated mostly with chemosensation and mechanosensation respectively (20, 40). This makes it difficult to disentangle the precise molecular mechanisms associated with the emergence of the predatory behaviour in *P. pacificus* using these mutants alone. However, by focusing on the role of mechanosensation, we have been able to identify *Ppa-mec-6* as a specific molecular component that has acquired additional functions in *P. pacificus* associated with its predatory feeding. In *C. elegans, Cel-mec-6* is a known to interact with several DEG/EnAC type subunits with the best characterised being *Cel-mec-4* and *Cel-mec-10* which form a mechanosensory ion channel responsible for the gentle touch response (24, 29, 52). Within this complex *Cel-mec-6* is a small auxiliary protein that is not a pore-forming subunit itself, but is necessary for proper channel function. In *P. pacificus*, we find mutations in any of *Ppa-mec-4, Ppa-mec-10* or *Ppa-mec-6* result in mechanosensory defects, but it is only *Ppa-mec-6* that has additional predation-specific abnormalities. Thus, while *Ppa-mec-6* may interact with *Ppa-mec-4* and *Ppa-mec-10* to regulate mechanosensation as in *C. elegans*, it is also likely involved with other as yet unknown components and may form a distinct mechanosensory channel involved in prey detection. These components require further identification.

By analysing the localization of *Ppa-mec-6*, we observed robust expression in the six head sensory IL2 neurons where it is also expressed in *C. elegans* (21) . These neurons are easily identified in *P. pacificus* by their morphology and soma location, which also show a distinct placement compared to *C. elegans* (44, 53). In both species, the IL2 neurons project neurites toward the nematode nose with the sensory endings exposed to the external environment. In *C. elegans*, these neurons regulate dauer formation and nictation behaviours, and are thought to be both chemosensory and mechanosensory (54, 55). Furthermore, they release extracellular vesicles potentially involved in communication (56, 57). In *P. pacificus*, our results show that IL2 inhibition also prevents the acquisition of sensory information necessary for prey detection. Crucially, recent studies in *P. pacificus* identified the IL2 neurons as regulators of predatory feeding states which depend on the balancing actions of octopamine and tyramine (17). Octopamine induces predatory states while tyramine initiates more docile bouts and IL2 neurons express octopamine receptors necessary for the predatory states in *P. pacificus*. Accordingly, our findings that these neurons express *Ppa-mec-6* as well as *Ppa-daf-19* reinforces their importance as prime candidates for detecting and determining prey contact events. Moreover, the convergence of both mechanosensory and chemosensory inputs, as well as behavioural state modulatory mechanisms, in the *P. pacificus* IL2 neurons demonstrates that even relatively simple circuits can be remodelled to accommodate complex and dynamic behaviours including predation. Therefore, taken together, our findings provide new insights into how sensory systems are co-opted and refined to generate behavioural diversity, and demonstrate the importance of sensory perception and evolutionary adaptations for establishing novel behavioural traits.

## Materials and methods

### Worm maintenance

*P. pacificus* (PS312 and derived strains) or *C. elegans* (N2), were maintained on nematodes growth media (NGM) 6 cm plates seeded with 300 μl *Escherichia coli* (OP50) at 20^°^C. All *P. pacificus* strains used are listed in supplementary table 1A. The *C. elegans* N2 strain was provided by the CGC, which is funded by NIH Office of Research Infrastructure Programs (P40 OD010440).

### Orthologue identification

We generated a mechanotransduction channel list from wormbook (22) and obtained published genome assemblies for eight nematode species belonging to different clades identified by Blaxter et al (23). These species include: *Pristionchus pacificus* (10), *Caenorhabditis elegans* (release WS271, 2019, WormBase website (58)), *Strongyloides ratti* (59), *Bursaphelenchus xylophilus* (60), *Brugia malayi* (61, 62), *Ascaris suum* (63), *Romanomermis culicivorax* (64) and *Trichinella spiralis* (65). When multiple annotated isoforms were available, only one was considered for analysis. Orthologous were determined using OrthoFinder (66).

### Generation of new strains

Mutations were induced by CRISPR/Cas9 (IDT) by co-injection of the sgRNA for the gene of interest while using a sgRNA targeting *Ppa-prl-1* as a marker as previously described (67). sgRNAs were made by mixing crRNA (IDT) and tracrRNA (IDT) in equal volume and quantity and incubated for 5 min at 95^°^C. After letting them cooldown 5 min at room temperature, up to three sgRNA were mixed with the Cas9 and 5 min later Tris-EDTA (TE, Promega) was supplemented to reach the concentration of 18.1 μM for sgRNA and 12.5 μM for Cas9. This mix was centrifuged for 10 min at maximum speed at 4^°^C. The resulting mix was used to inject *P. pacificus* adults in the gonads area (Axiovert 200, zeiss and Eppendorf set up). P0 injected worms were singled out and laid eggs overnight. Once the progeny had developed enough, plates were screened for a roller phenotype induced by the dominant mutation in *Ppa-prl-1* pointing to successful injection. Around 50 worms were genotype from identified plates for the gene of interest using PCR (Qiagen) followed by sanger sequencing (Eurofins). The sgRNA and primer sequences used as well as the wild-type sequences and generated mutation are provided in supplementary table 1B. Two guide RNA were used for *Ppa-mec-3* to target both isoforms. All mutations generated are predicted to lead to an early stop codon or frameshift with the exception of one allele of *Ppa-mec-3 (bnn61)* and one allele of *Ppa-trp-2 (bnn51*) presented in supplementary figures. Mutations were generated in the *P. pacificus* PS312 strain.

Behavioural tracking and state predictions required an integrated *Ppa-myo-2p*::RFP (17). In addition, mouth morph ratio is influenced by *Ppa-daf-19* which results in a strong st bias unsuitable for predation assays (18). Therefore, predatory morphs eu were induced alongside *Ppa-daf-19* through generation of additional mutations in *Ppa-nag-1; Ppa-nag-2* (18) in the *Ppa-myo-2p*::RFP background. Accordingly, this strain JWL118 has a 100% Eu mouth morphs (Supplementary Fig. 3A). For experiments involving *Ppa*-*daf-19*, all comparative experimental strains also carry *Ppa-nag-1; Ppa-nag-2* mutations based on the control strain JWL147 (see supplementary table 1).

### Generation of reporter lines

TurboRFP and GFP sequences optimized for *P. pacificus* were retrieved from plasmid pZH009 and pZH008 respectively (9). A Venus reporter was optimised for *P. pacificus* as previously described (9). Promotor sequences consisted of the indicated number of nucleotides preceding the ATG according to *P. pacificus* genome (10): *Ppa-mec-6* (949 bp), *Ppa-mec-3* (932 bp), *Ppa-daf-19* (719 bp) promotor sequences. *Ppa-daf-19p*::RFP (pMR027, Genscript) and *Ppa-mec-6p*::Venus (pMR022, eurofins) were made by gene synthesis and *Ppa-mec-3p*::RFP (pMR005) by cloning. For generating stable transgenic lines, animals were injected with a mix containing 60 ng/μl purified gDNA (NEB Monarch purification kit) along with 5-10 ng/μl plasmid of interest all digested with HindIII. Prior to each injection, the mix was centrifuged at maximum speed at 4^°^C for 10 min. Injection were made in *P. pacificus* adult gonads (Axiovert 200, zeiss, and Eppendorf set up). Progeny worms carrying the construct of interest were identified using the epi-fluorescence microscope (Axio Zoom V16; Zeiss). Confocal images were acquired using a stellaris microscope (Leica) with a 63x objective at water immersion.

### Chemosensory assays

Attraction toward OP50 was tested using previously developed assays with minor modifications (40). 20 μl of an OP50 overnight culture was pipetted onto a 6 cm unseeded plate 1.5 cm from the edge and left at room temperature overnight. Plates where then kept in the fridge until use. 50 adult worms from plates almost depleted of food were placed at the opposite side of the bacterial lawn at around 1.5 cm of the border making sure to not transfer any bacteria when doing so. After 8 hours the number of worms at the bacterial lawn and the total number of worms still alive in the plates were determined. Plates were scored if they had at least 30 worms left at the end of the assay.

To assess worm behaviour toward 1-octanol, we adapted existing protocols (41). Adult worms were washed off with M9 and left on an unseeded plate for 3 hours to remove bacteria. Prior to starting the assay, two drop of 1.2 μl of sodium azide 1M was deposit 3 cm apart on opposite side of a 6 cm unseeded plate. 100 worms were placed in the middle line between the two drops and immediately 1μL of 100% ethanol was put as control on top of one of the sodium azide deposits and 1 μl of 100% 1-octanol (sigma-aldrich) over the other. The assay plates were left for 16 hours at 20^°^C before assessing worm position. Only worms within a 2 cm radius of the drop up to the midline were taking into account leading to a total of at least 40 worms. Chemosensation index was calculated as the ratio between the number of worm closer to the 1-octanol minus those closer to the ethanol relative to the total number of worms. For both assays 10 replica were made over at least three different days.

### Touch assays

Touch assays were performed as described for *C. elegans* with minor modifications (22). Worms were transferred to unseeded plates and left to recover for several minutes prior to being utilised in one of the three assays. Harsh touch assays consisted of touching the worm between its vulva and its tails with the platinum pick used in every day worm maintenance. For the gentle touch assays, the worm was caressed with an eyelash (taped to a wood skewer pick for handling) between its vulva and its tails. For both assays, each worm was tested ten times and the number of times that trigger a reaction was assessed. Due to the fact that some of our strains appear lethargic we did not only consider forward movement as a reaction but any change in worm state occurring upon contact. To assess sensation of the nose, the eyelash was presented to the nose of the worm ten time and again all sign of detection were considered (backward movement or head bending). At least 30 worms over at least 3 different days were assessed for each test and strain.

### Corpse assay

Predatory behaviour was tested as previously described with modifications (11, 17). *C. elegans* larvae were collected from plates depleted in food just after hatching using M9 and passed through 20 µm filter to remove the other developmental stages. After centrifugation (2000 rpm, 1 min), ∼1 μl of larvae was put off centre of a 6 cm unseeded plate, see supplementary figure 1A. In parallel, predators consisting of young adult *P. pacificus* of the strain of interest were picked onto another unseeded plates. Both predators and prey were left for 2 hours to allow the larval prey to spread and predator to starve. 5 predators were added on the plates and a copper arena 2.25 cm^2^ was put around them. Assays were left for 2 hours then two pictures of the arena were taken at ∼20x (each field covered a quarter of the arena) at least 10 second apart (Axio Zoom V16; Zeiss). The two images were process with Fiji using the ‘max’ set up and corpses were manually count on the resulting image. To assess kin recognition PS312 *P. pacificus* larvae were used as prey with 20 predator and the assay took place without the arena for 24 hours. Corpses were counted under the bench microscope (stemi 508, zeiss). For both assay 10 replicates were conducted at least three different days. To increase the power of our assay and detect smaller variation between strain, 20 replicas were made for figure 3.

### Mouth form assessment

Worms were grown in OP50 plates until a high number of adults was reach. Worms were then washed in M9 and placed on a slide containing sodium azide to immobilized them. The number of eu and st worms were counted under an Axiovert 200 (zeiss) at 10x or 20x. Mouth form was controlled from 5 different plates over at least three different days with a minimum of 30 animals scored per assay.

### Feeding assay with PharaGlow

PharaGlow experiments were performed as described previously (17). Briefly, adult worms with a fluorescent pharynx marker (*Ppa-myo-2*p::RFP) were picked onto an unseeded plates and left for a 2 hour starvation period. 30 worms were picked onto unseeded assay plates containing *C. elegans* larvae made as for the killing assays (see above) but with twice the density to saturate the plate in larvae. To maintain the worms inside the field of view, a copper arena 2.25 cm^2^ was put around the assay animals. Worms were acclimated to their new environment for 15 min before recording. Recording was made with an epi-fluorescence microscope (Axio Zoom V16, Zeiss) at 1x effective magnification (60 N-C ⅔’’ 0.5 x, Zeiss) and Basler camera (acA3088-57um, BASLER). Light intensity was set at 100% and acquisition time was adjusted to use the full range of intensity without saturating the signal. Images were acquired at 30 Hz for 10 min. Images were then compressed and processed with the python package PharaGlow (https://github.com/scholz-lab/PharaGlow), (43) adapted for *P. pacificus* (17) with the following parameters (subtract:1, smooth:3, dilate:3, tfactor:0.9, thresholdWindow:50, bgWindow:500, length:70, watershed:100, minSize:250, maxSize:1000, searchRange:10, memory:30, minimalDuration:1800, widthStraight:10, pad:5, nPts:50, linewidth:2). The quality of extracted movies was manually assessed and any worms with no sign of activity throughout the recording length were considered dead or damaged and removed from the analysis. Post-processing to calculate the velocity and pumping rate was done using a previously developed python script and finally automatic behavioural prediction were assigned using the git-hub package PpaPred as described previously (https://github.com/scholz-lab/PpaPred), (17). Worms were recorded over at least three different days.

### Inhibition of neurons by histamine chloride channels

The *Ppa-mec-6* promotor from our reporter construct *Ppa-mec-6p*::Venus (pMR022) was cloned in front of the *P. pacificus* codon optimized histamine chloride channel from plasmid pLJ149 (17) forming plasmid pLB010. Transgenic line JWL237 carrying both pMR022 and pLB010 was generated as described above in the wildtype PS312 background. JWL237 was then crossed in the *Ppa-myo-2p*::RFP background (JWL27) for tracking experiments. Only worms displaying green fluorescent were used for experiments.

Histamine plates were made by the addition of histamine chloride to the NGM before pouring to reach a final concentration of 10 mM. Plates were kept and used in the dark to preserve the histamine. Touch, chemosensation and feeding assays were made as described above with worm transferred to the histamine plates at least 10 minutes before performing the assay or from the starvation period when applicable. For corpses assays, copper arenas were not used as they reacted with the histamine chloride and due to the wide field of exploration, images could not be taken and instead corpses were counted manually after 2 h exposure to predators. For developmental assay and mouth form assessment, histamine plates were seeded as usual with 300 μl of OP50 and around 100 J2 larvae were added following filtration (20 µm filter, Millipore). After 4 days at 20^°^C, the number of eu or st animals were assessed.

### Data representation and statistical analysis

Box plot illustrates data distribution with quartiles Q1, Q2, and Q3 representing the 25th, 50th, and 75th percentiles, respectively. The box, bounded by Q1 and Q3, includes a thicker line at the median (Q2). Whiskers show the range where most values fall. Analysis of chemosensory assay, touch assay, corpses assays and mouth form ratio was made with R software and its package prettyR with significance determine using one-direction Wilcoxon Mann Whitney with Benjamini-Hochberg correction (68). Analysis of the state prediction results (relative time in state, mean bout duration, transition rates) were done with python as describe previously (17) relying on Wilcoxon Mann Whitney with Bonferroni correction. In figures, statistics results are display as follow: non-significant (ns), p-value ≤ 0.05 (*), ≤ 0.01 (**), ≤ 0.001 (***), ≤ 0.0001 (****).

## Acknowledgments

We would like to thank all the Lightfoot and Scholz labs for useful discussions, Desiree Goetting for critical reading of the manuscript, Wolfgang Bönigk for plasmid cloning (Genetics Facility, MPI for Neurobiology of Behavior-caesar), the Imaging facility for assistance (MPI for Neurobiology of Behavior-caesar) and Jürgen Berger for SEM imaging presented in figure 5 (MPI for Biology). The strains PS312 and RS3238 were provided by the Sommer lab (MPI for Biology, Tübingen).

