## Supplementary figures and images for "Evolution of sensory systems underlies the emergence of predatory feeding behaviours in nematodes"

### Supplemental Files

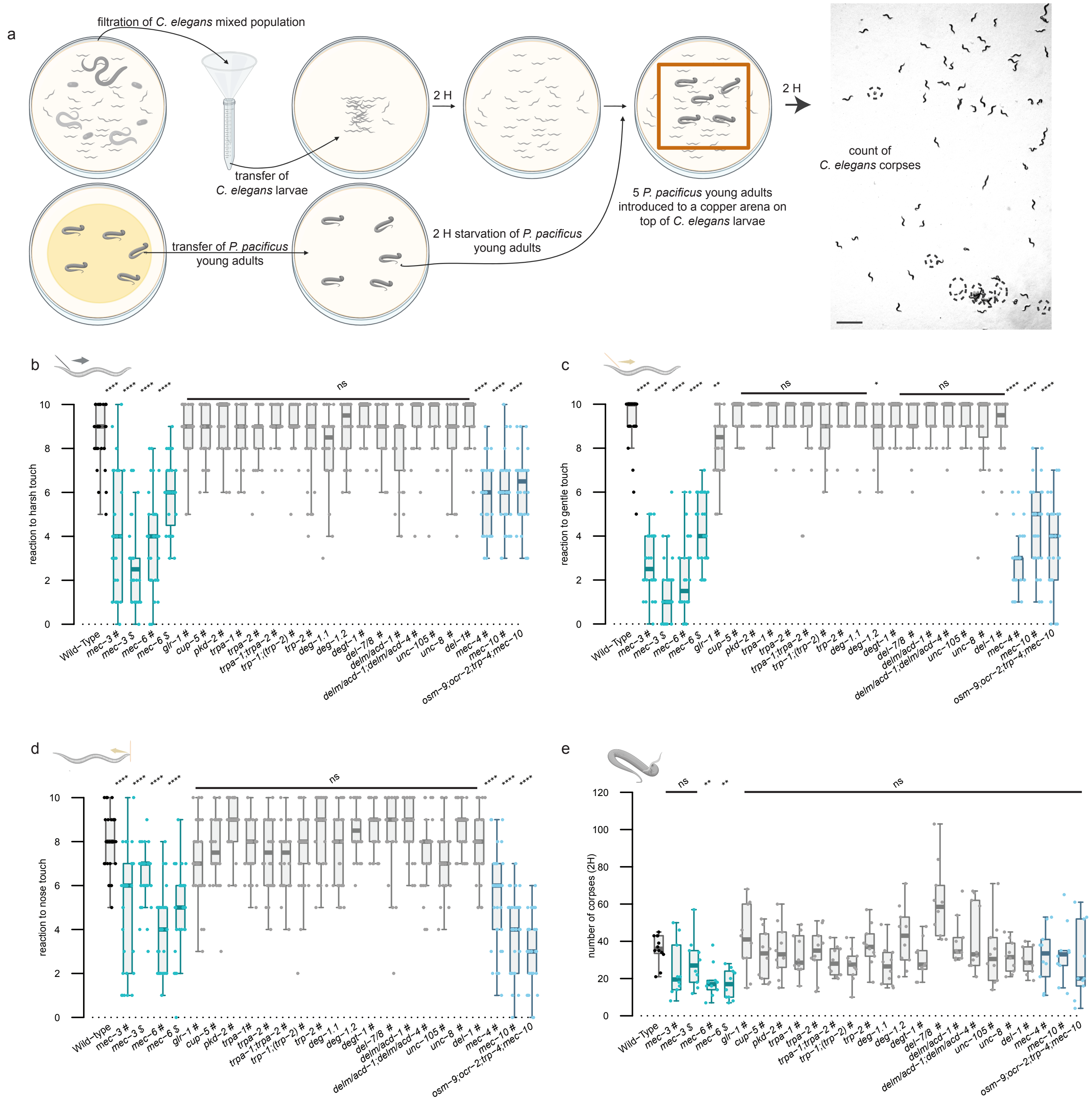

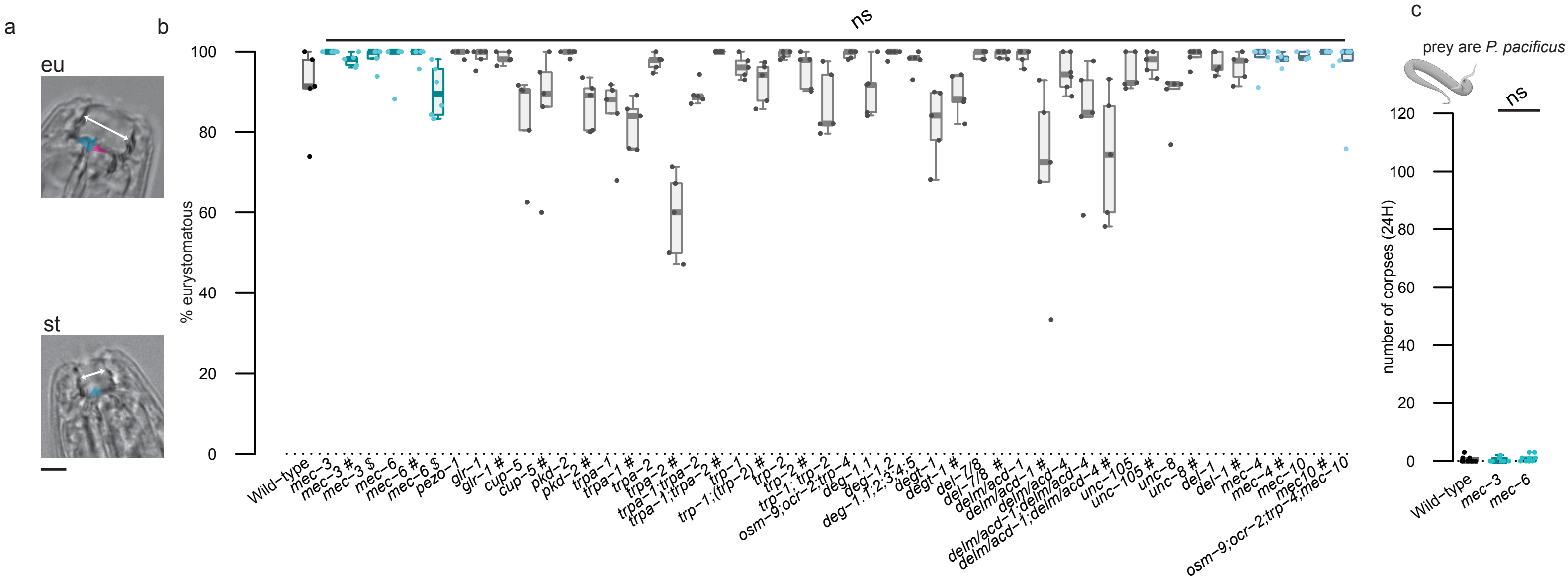

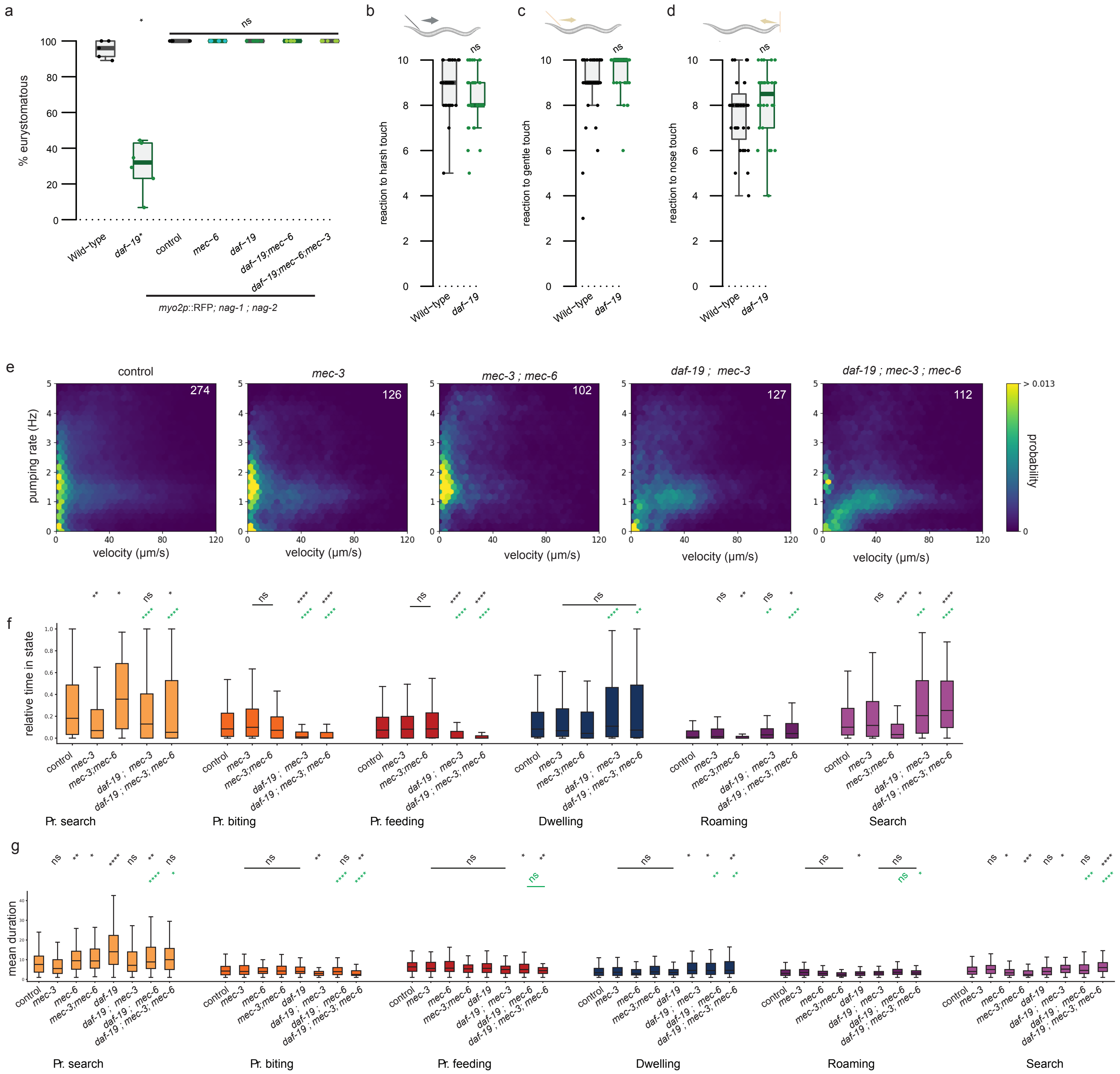

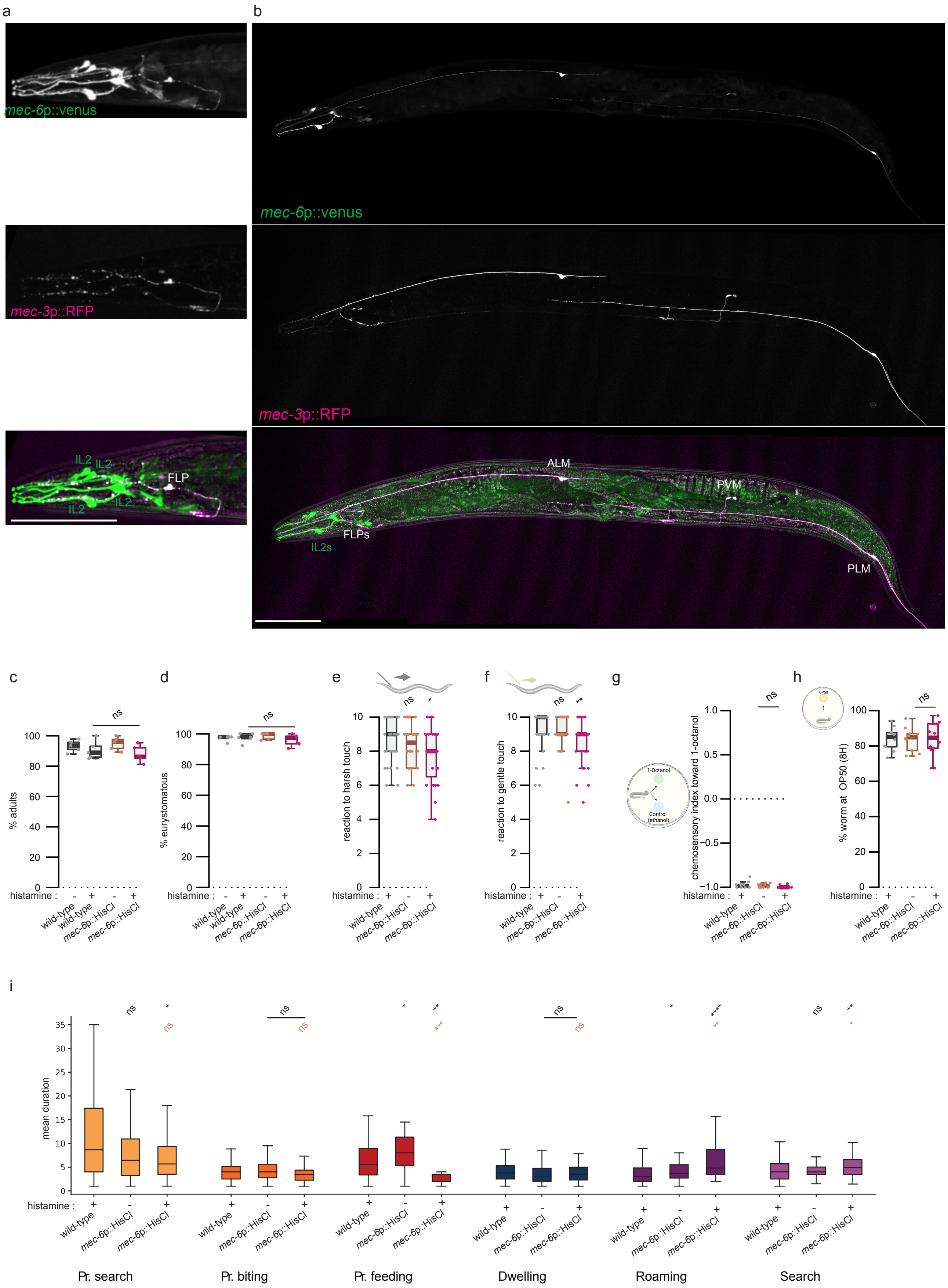
